# Hippocampal sclerosis of aging at post-mortem is evident on MRI more than a decade prior

**DOI:** 10.1101/2023.03.08.531683

**Authors:** Diana Ortega-Cruz, Juan Eugenio Iglesias, Alberto Rabano, Bryan Strange

## Abstract

**INTRODUCTION:** Hippocampal sclerosis of aging (HS) is an important component of combined dementia neuropathology. However, the temporal evolution of its histologically-defined features is unknown. We investigated pre-mortem longitudinal hippocampal atrophy associated with HS, as well as with other dementia-associated pathologies.

**METHODS:** We analyzed hippocampal volumes from MRI segmentations in 64 dementia patients with longitudinal MRI follow-up and post-mortem neuropathological evaluation, including HS assessment in the hippocampal head and body.

**RESULTS:** Significant HS-associated hippocampal volume changes were observed thoughout the evaluated timespan, up to 11.75 years before death. These changes were independent of age and Alzheimer’s Disease (AD) burden, and specifically driven by CA1 and subiculum. AD burden, but not HS, significantly associated with the rate of hippocampal atrophy.

**DISCUSSION:** HS-associated volume changes are detectable on MRI earlier than 10 years before death. These findings could contribute to the derivation of volumetric cut-offs for *in vivo* differentiation between HS and AD.

## 1. Introduction

Hippocampal sclerosis of aging (HS) is a pathological finding associated with worsened cognitive symptoms in dementia patients [1,2]. Since its first description by Dickson *et al*. [3], it has been defined by a severe neuronal loss in CA1 and subiculum subfields of the hippocampus, which is particularly prevalent in patients over 85 years of age [4]. This qualitatively-observed lesion is disproportionate to expected damage by other coexisting pathologies. HS is normally found in combination with one or several entities across the dementia neuropathological spectrum: Lewy bodies (LB) [5,6], vascular pathology [7,8], Alzheimer’s disease (AD) [9,10], and most commonly, limbic-predominant age-related TDP-43 encephalopathy (LATE) [11,12]. Indeed, 80-100% patients with HS present LATE pathology, and HS is found in up to 90% of LATE cases [13].

Despite its relevance within the neuropathological spectrum of dementia, the contribution of HS is often overlooked in the field. Several recent post-mortem studies on AD [14,15], TDP-43 [16,17] and LB pathologies [18,19] do not include HS within their evaluation. Conversely, other studies have highlighted the importance of HS in dementia [20,21], which allowed the observation of histological findings proposed as early HS [5,22]. In line with this work, we recently characterized early pathological changes of HS, proposing this pathology forms a wider spectrum than initially defined [23]. Following this re-classification, the prevalence of HS could have been underestimated in previous dementia studies, which reported a frequency of 3-30% following the classical definition [1,2,9].

The definition of HS by histological findings has limited the search for biomarkers of this pathology. Currently, HS can only be diagnosed post-mortem [24]. Preliminary evidence suggests that patients with HS have lower Mini-Mental State Examination (MMSE) scores and longer symptom duration than AD patients without HS [10], showing potential as a clinical biomarker. HS patients also show relative preservation of verbal, executive and visuomotor functions compared to AD [25,26]. Moreover, HS has been associated with end-stage reduction of hippocampal volume detectable in pre-mortem MRI [27]. However, the identification of these changes as tentative biomarkers is limited by a current lack of understanding of the timespan of HS. Since this pathology can up to now only be evaluated post-mortem, the onset and duration of its histological hallmarks remains unknown.

In this study, we aimed to characterize the timespan of hippocampal volume changes associated with HS. We studied data from dementia patients with pre-mortem MRI follow-up, spanning more than 10 years before death, and subsequent neuropathological evaluation. We describe longitudinal differences of whole hippocampus and subfield volumes between subjects with HS (HS+) and without (HS-), following its classical definition, as well as according to our recently proposed evaluation including early stages [23]. Implementing a two-level HS assessment in the head and body of the hippocampus, we explore subfield volumetric changes in these two regions relative to HS. We also evaluate longitudinal volume differences as a function of other neuropathologies, thus dissociating gradients associated with each pathology from those of HS – which is crucial for differentiating HS from AD with *in vivo* imaging.

## 2. Methods

### 2.1 Cohort

We studied data obtained between 2007 and 2020 from the Vallecas Alzheimer’s Center Study (VACS). This center is a nursing home for dementia patients who undergo clinical and neuroimaging follow-up. The follow-up protocol includes baseline and semestral assessments. We included patients with at least one pre-mortem MRI acquisition and donation to the BT-CIEN brain bank after passing away, allowing neuropathological evaluation. Out of 102 patients meeting these criteria, 10 were discarded due to the following exclusion criteria: a) no T1 scan without considerable movement artifact (n=4); b) a neuropathological diagnosis of Frontotemporal Dementia (n=3), given its independent association with hippocampal sclerosis [9]; c) unattainable evaluation of hippocampus slides due to extreme MTL atrophy (n=1) or hemorrhage (n=2).

### 2.2 Neuropathological evaluation

Brain extraction and processing followed previously described BT-CIEN procedures [28]. The post-mortem interval was kept as short as possible, with an average duration of 4.4±1.9h. After extraction, the left hemisphere was fixed for neuropathological diagnosis and cut into coronal slices for tissue block dissection. Blocks were embedded and cut into 4µm sections for hematoxylin/eosin and immunohistochemical staining (against amyloid β, phospho-tau AT100, total α-synuclein, and total TDP-43).

Neuropathological evaluation was performed following published consensus criteria, and patients were stratified into low- and high-burden groups for each pathology. Probability (0-3) of Alzheimer’s Disease Neuropathological Change (ADNC) was assessed based on National Institute on Aging (NIA) guidelines [29], including Thal Aβ stage, Braak stage, and neuritic plaque score. Subjects with a high ADNC probability were classified into the high AD burden group. Cerebrovascular pathology was evaluated following the staging defined by Deramecourt *et al*. [30], which includes partial scoring in the frontal and temporal lobes, basal ganglia and hippocampus, added into a total vascular score (0-20). We classified patients with a total score of 8 or higher as having high vascular burden. Lewy body pathology was assessed through Braak α-synuclein staging (0-6) [31], and high burden was assigned to those with a value higher than 2 (pathology beyond the brainstem). TDP-43 proteinopathy was evaluated according to LATE staging (0-3) [11], and patients with inclusions in the hippocampus (stage 2 or higher) were categorized as high LATE burden. HS was assessed following our recently proposed staging system (0-IV) including early stages of the pathology [23], evaluated in sections of both head and body of the hippocampus. Subjects presenting stages>0 in any of these two sections were classified as HS+. Its classical definition was also employed for alternative HS classification [4], with subjects presenting severe neuronal loss in CA1 and subiculum (stage>II) of hippocampal head sections classified as HS+.

### 2.3 MRI acquisition

MRI scans were acquired using a 3T scanner (Signa HDxt General Electric, Waukesha, USA), and with a phased array eight-channel head coil. T1-weighted images were obtained using a 3D sagittal sequence Fast Spoiled Gradient Recalled (FSPGR) configuration with inversion recovery (repetition/echo/inversion times 7/3.2/750ms, field-of-view 240mm, matrix 288×288, slice thickness 1mm), yielding a resolution of 1×0.469×0.469mm. The quality of acquired scans was evaluated by an experienced radiographer.

### 2.4 Longitudinal hippocampal segmentation

Volumes of the whole hippocampus and its subfields were obtained by automated segmentation in FreeSurfer 7.1.1. For that purpose, all T1-weighted scans available from subjects included in the study were processed with the FreeSurfer longitudinal processing workflow [32]. Since a considerable proportion of scans presented slight movement artifacts, subcortical segmentations were substituted by the segmentation from *SynthSeg* [33,34] to increase the accuracy of the subsequent step. Segmentation of hippocampal subfields and amygdaloid nuclei [35,36] was then obtained and visually inspected to discard those with inaccurate subfield delineation. Out of 92 subjects, 7 resulted in errors along the processing pipeline and 21 had segmentations from all scans discarded, resulting in a total of 64 subjects with accurate volumetric data. Left and right hemisphere volumes were averaged and used for subsequent statistical analyses.

### 2.5 Statistics

RStudio 11.4.1106 software was used for statistical analyses and plots, with packages *lme4, emmeans, tidyverse*, and *ggpubr*. To compare between HS groups, for categorical, non-normal numeric, and normal numeric variables, we used Pearson’s chi square, Kruskal-Wallis and t-tests respectively. Linear mixed models with random intercept and slope were employed to study volume changes of the hippocampus or subfield of interest. These dependent variables were analyzed in each model as a function of time, assessing its interaction with the pathology of interest. In addition, age at death, sex and intracranial volume were included as independent variables in the model. Pathological groups were represented by factor variables, while HS stages were coded as ordinal variables.

Variable effects were assessed by analysis of variance and estimated marginal means at different timepoints were compared between groups.

### 2.6 Ethical approval

The BT-CIEN brain bank is officially registered by the Carlos III Research Institute (Ref: 741), by which donation is carried out under informed consent by a relative or proxy. BT-CIEN procedures and VACS follow-up have been approved by local health authorities (Ref: MCB/RMSFC, Expte: 12672). The study of this data has been independently approved by the Ethics Committee of the Universidad Politécnica de Madrid (Nº Expte: 2021-062).

## 3. Results

### 3.1 Cohort features

Demographic and follow-up details, as well as neuropathological burden distribution from the 64 patients with accurate hippocampal segmentations are shown in Table 1. Variables are also compared between HS- and HS+ patients, following the staging system including early stages of the pathology. The studied cohort is composed by a predominantly female population of advanced age at death. HS+ subjects displayed expected differences associated with the pathology, with significantly longer disease duration (reflected by time in nursing home), older age, and lower final severe-MMSE scores. As expected, this group also presented higher burdens of ADNC and LATE pathology. Results were not altered if patients were classified following the classical HS definition, including only advanced stages.

**Table 1.** Demographic, follow-up and neuropathological data for subjects included in volume analyses in this study. NOTE. The three left-most columns show values for groups with and without HS, with early and advanced stages included within the HS+ group. Results for comparison between groups do not change when stratifying groups by the classical HS definition. Group comparisons by: ^a^ Pearson’s chi-2 test. _b_ Kruskal-Wallis test. ^c^ T-test. *Age at symptom onset is estimated by neurologists based on medical records and interviews with caretakers. Abbreviations: ADNC: Alzheimer’s Disease Neuropathological Change; HS: hippocampal sclerosis of aging; LATE: limbic age-related TDP-43 encephalopathy; MRI: Magnetic Resonance Imaging; SD: standard deviation; sMMSE: severe Mini-Mental State Examination; TDP-34: TAR DNA-binding protein 43.

|  | Whole cohort | HS- (n=26) | HS+ (n=38) | p value |
| --- | --- | --- | --- | --- |
| **Demographics** | Mean ± SD or % | | | |
| Sex (%Females) | 81.3 | 73.1 | 86.8 | .166<sup>(a)</sup> |
| Estimated age at symptom onset * | 76.5 ± 6 | 75 ± 6.5 | 77.6 ± 5.5 | .354<sup>(b)</sup> |
| Age at death | 88.4 ± 6.6 | 84.6 ± 6.6 | 90.9 ± 5.3 | **10<sup>-4</sup>**<sup>(c)</sup> |
| sMMSE score at admission | 19.1 ± 9.2 | 20.5 ± 9.3 | 18.3 ± 9.1 | .287<sup>(b)</sup> |
| sMMSE score prior to death | 10.4 ± 10.6 | 14.9 ± 11 | 7.4 ± 9.3 | **.005**<sup>(b)</sup> |
| **Follow-up** | Mean ± SD | | | |
| Time in nursing home (years) | 4.7 ± 3.4 | 3.3 ± 3.3 | 5.7 ± 3.2 | **.004**<sup>(b)</sup> |
| Time between last MRI and death (years) | 3 ± 3.1 | 1.6 ± 2.2 | 4 ± 3.3 | **4·10<sup>-4</sup>**<sup>(b)</sup> |
| Number of MRI scans included | 3.2 ± 2.7 | 3 ± 2.5 | 3.3 ± 2.9 | .849<sup>(b)</sup> |
| **Neuropathology burden** | % |  |  |  |
| High ADNC | 68.8 | 46.2 | 84.2 | **.001**<sup>(a)</sup> |
| High vascular pathology | 60.9 | 73.1 | 52.6 | .1<sup>(a)</sup> |
| Lewy bodies beyond brainstem (Braak>2) | 40.6 | 34.6 | 44.7 | .418<sup>(a)</sup> |
| TDP-43 inclusions in hippocampus (LATE>1) | 50 | 11.5 | 76.3 | **6·10<sup>-7</sup>**<sup>(a)</sup> |
| Early or advanced HS | 59.4 | 0 | 100 | - |
| Classical HS | 29.7 | 0 | 50 | **2·10<sup>-5</sup>**<sup>(a)</sup> |

Included scans from pre-mortem MRI follow-up ranged between 1 and 11 per subject and covered a timespan of up to 11.75 years before death. Time between last included MRI and death was also significantly higher in the HS+ group, due to the feasibility of correct MRI acquisition being directly affected by the cognitive state of the patient.

### 3.2 Longitudinal hippocampal atrophy as a function of HS

Following the two HS classification criteria evaluated here, we compared changes in whole hippocampal volumes between HS- and HS+ subjects across time. Given the association of HS with age, we included age at death in the model, as well as sex and intracranial volume. Including early and advanced stages, patients with HS displayed reduced hippocampal volumes compared to the HS-group (F=31.7, p=.006), as shown in Figure 1A. This difference was significant throughout the whole evaluated timespan, based on comparison of estimated marginal means (EMM) at each year. At 11.75 years before death, the EMM difference between HS- and HS+ subjects was 405±146 mm^3^ (T=2.8, p=.007). Critically, the rate of decline in volume was not significantly different between groups (estimated slopes (mm^3^/year): -34.4 for HS+, -43.9 for HS-). Significantly lower volumes without different decline rates were also found following the classical HS definition (Figure 1B), showing this association is independent of the classification criteria (F=25.6, p=.005; HS+ slope=-41.9. HS-slope=-36.4). Volume effects of HS with these two classifications were still significant in a subset of the cohort including 26 HS+ subjects with closest ages to HS-ones, further supporting the independence of results on age (mean age at death for this HS+ subset: 88.1±3). These effects were also independent of AD pathology burden, as adding this variable in the model did not remove the significant effect of HS throughout this timespan (Figure S1).

**Figure 1.**
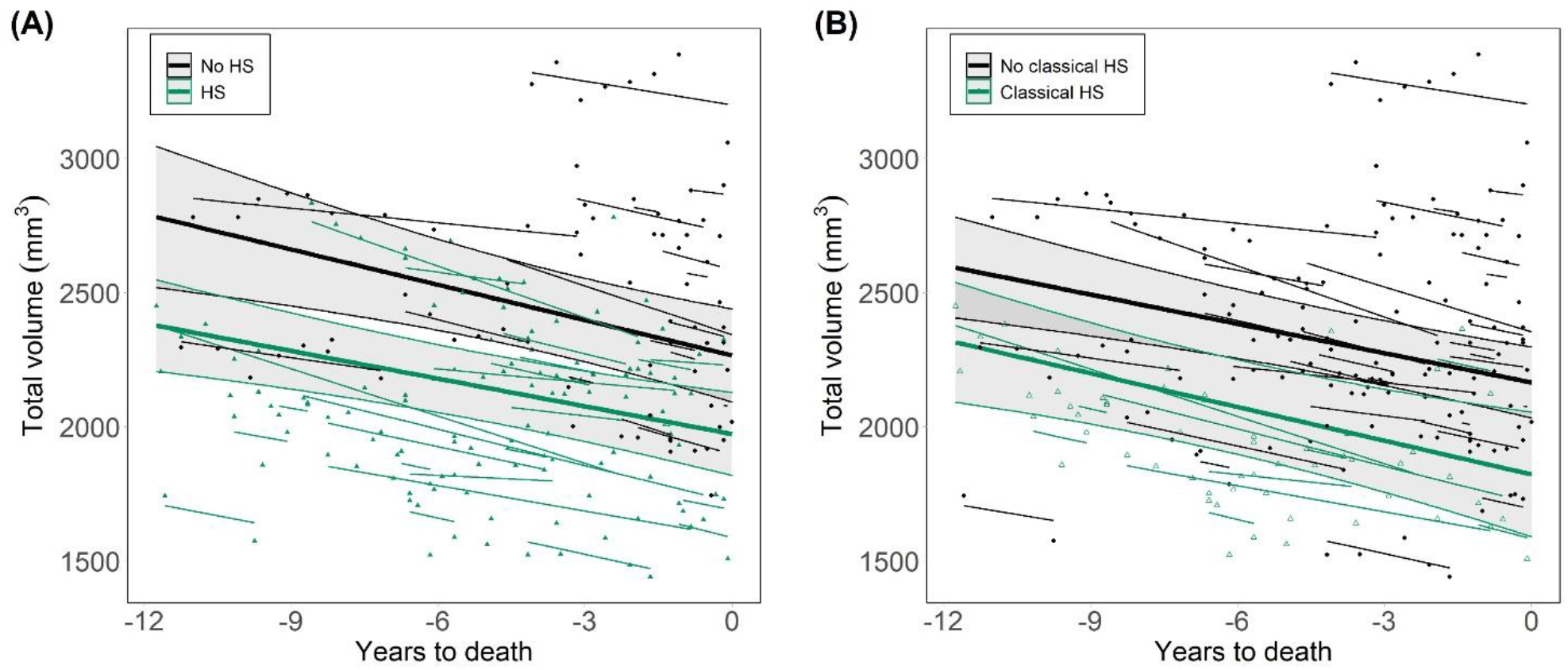
Longitudinal progression of hippocampal volumes as a function of HS. (A) HS groups defined as a function of HS staging including early and advanced stages (HS+ for those at stage>0) [23]. (B) HS groups determined by the classical definition of the pathology, by which only advanced cases with severe cell loss are included within the HS+ group. HS: hippocampal sclerosis of aging.

### 3.3 Hippocampal subfield volume changes relative to HS staging

To test the regional specificity of the observed effects, we analyzed the association between HS stage and subfield volumes in both head and body of the hippocampus. Hippocampal subfields were analyzed in two groups: CA1+subiculum, in which histological changes occur at earlier HS stages, and CA3+CA4, which may present changes in stage IV [23]. Through the evaluated timespan, volumes of CA1+subiculum were significantly lower in both hippocampal head (Figure 2A) and body (Figure 2B) as a function of respective HS stages (head: F=5.8, p=.035; body: F=8, p=.003). Such association was not observed for CA3+4 volumes in the head (Figure 2C) or body (Figure 2D) of the hippocampus. Rates of change in subfield volumes were not significantly different as a function of HS staging, nor between HS+ and HS-groups. We also evaluated volume changes in CA1 and subiculum separately as a function of HS (Figure S2), comparing EMM to test for group differences through the timespan. CA1 in the hippocampal head and subiculum in the body were the two subfields displaying significant group differences, with lower volumes in HS+ as early as 11 and 9.5 years before death, respectively. These reported results as a function of HS classification including early stages were also significant following the classical HS definition.

**Figure 2.**
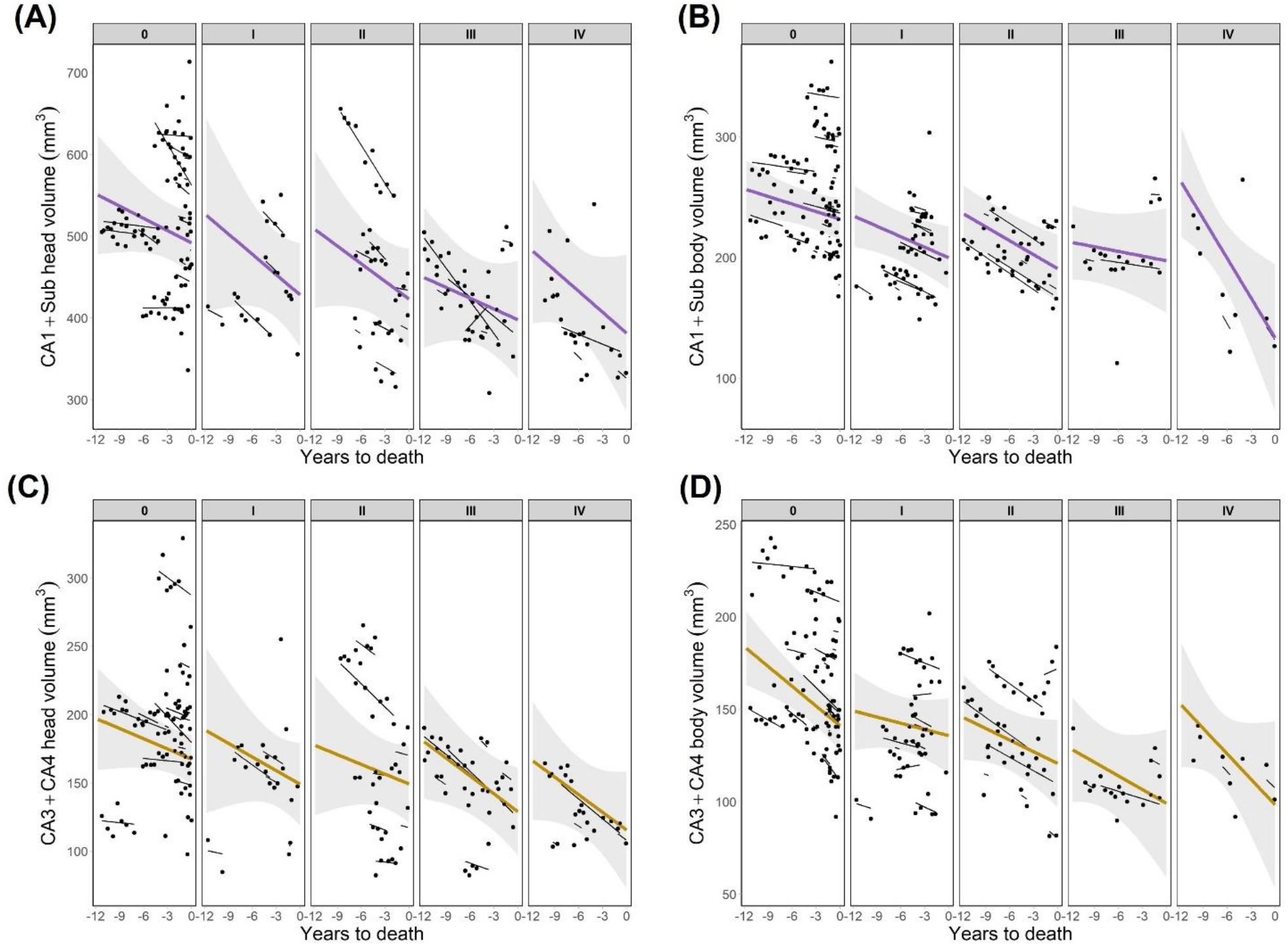
Longitudinal progression of hippocampal subfields volumes as a function of HS staging. (A) Volumes of CA1+subiculum in the head of the hippocampus. (B) Volumes of CA1+subiculum in the hippocampal body. (C) CA3+CA4 volumes in the head of the hippocampus. (D) CA3+CA4 volumes in the hippocampal body. CA1+Sub: CA1+subiculum.

### 3.4 Longitudinal hippocampal atrophy as a function of other pathologies

Given that we did not find an effect of HS on the rates of hippocampal volume reduction, we evaluated whether any other neuropathology presented such association. Whole hippocampal volumes across time were separately analyzed as a function of AD, vascular, LB and TDP-43 pathologies. To enable group comparisons in this dementia cohort, patients were divided into high- and low-burden groups for each pathology. As shown in Figure 3, groups displaying high probability of ADNC (F=14.9, p=5·10^−4^) as well as high LB (F=4.6, p=.027) and TDP-43 burden (F=13.6, p=.015) showed significantly lower volumes compared to their respective low-burden groups. In addition, the effect of ADNC severity on hippocampal volume displayed a significant interaction with pre-mortem time, and thus different decline rates between groups (estimated slopes: -13.3 for low/intermediate, -44.8 for high ADNC). Considering baseline and final MMSE scores, cognitive decline followed a similar trend as hippocampal volume: HS+ subjects showed lower scores, with no difference in decline rates, while high ADNC was associated with lower scores and steeper decline rates (Figure S3). No significant volume differences were found between low and high vascular burden groups.

**Figure 3.**
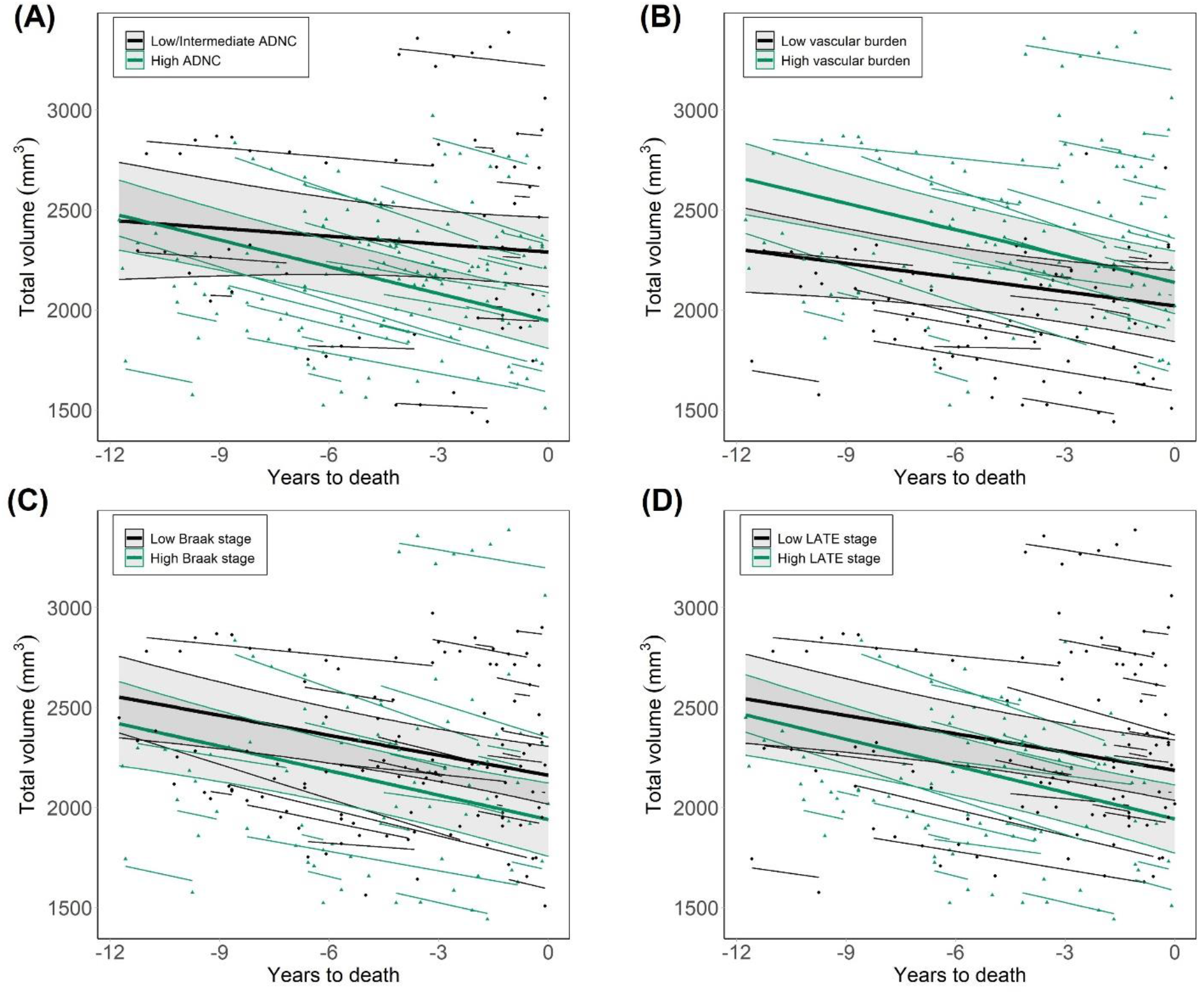
Longitudinal volumes of the whole hippocampus as a function of other neuropathologies in dementia. (A) Differences between groups of low/intermediate and high probability of ADNC. (B) Groups of low and high vascular pathology burden, following the evaluation proposed by Deramecourt *et al*. [30] Groups of low and high Lewy body pathology burden, as a function of Braak staging for α-synuclein. Comparisons between groups of low and high TDP-43 pathology burden, evaluated through LATE staging. ADNC: Alzheimer’s disease neuropathological change. LATE: limbic age-related TDP-43 encephalopathy.

## 4. Discussion

We studied longitudinal changes in hippocampal volume as a function of hippocampal sclerosis of aging, a histology-defined pathology for which no clinical or imaging biomarkers exist. Previous work by our group [23] and others [27,37] revealed an association between HS and hippocampal atrophy, with lower volumes in HS+ patients in all subfields of the hippocampal cortex. However, the temporal evolution and putative onset of these changes remained unknown. We found lower hippocampal volumes in HS+ compared to HS-subjects throughout the whole evaluated timespan, up to 11.75 years before death. The rate of volume change did not differ between groups, suggesting reduced volumes in HS+ patients manifest prior to this evaluated timespan. Importantly, this trend was consistent independently of age and AD burden. Our findings were also independent of the classification of early HS cases, indicating volume effects are driven by advanced stages, which present the classical HS signature defined by cell loss.

Hippocampal atrophy is a well-established marker of AD [38-40]. However, other neuropathologies have also been found to impact hippocampal volume, including vascular [41,42] and TDP-43 pathology [21,43], thus questioning the specificity of hippocampal volume as an index of ADNC. LB pathology is also associated with reduced hippocampal volumes [44], although less severely than in AD [45,46]. The rate of hippocampal atrophy has also been described as a useful predictor of AD [47-49], but this gradient had not been previously dissociated from other pathologies. Our results show that while ADNC, LB, LATE and HS all correlate with lower hippocampal volumes in symptomatic stages of dementia, its rate of change is only associated with ADNC burden. This suggests that, unlike other neuropathologies, AD continues to affect hippocampal atrophy beyond dementia onset. Thus, the decline in hippocampal volume across several timepoints could be a useful and specific AD biomarker, compared to single-timepoint atrophy measures.

Our subfield analyses revealed HS-associated hippocampal atrophy through the evaluated timespan is driven by lower CA1 and subiculum volumes, consistent with its histological pattern. More specifically, CA1 in the hippocampal head displayed this reduction earliest, and thus may serve as a useful target for early identification of HS. One distinguishing feature of our work is the histological evaluation of HS at two coronal levels of the hippocampal long axis [50], allowing the comparison of regional subfield volumes to their respective stage. In contrast, other studies assess HS in a single hippocampal section, either unspecified [4,7,20] or at the level of the lateral geniculate body [3,22], prompting variable results in the study of HS. The early reduction in CA1 head volume reported here points towards the anterior hippocampus as a key region in the pathology, and highlights the relevance of assessing HS in this level.

In addition to the two-level assessment of HS, allowing the evaluation of regional subfield associations, other strengths of this study include an extensive neuropathological evaluation and unified MRI acquisition and processing pipelines. Moreover, segmentation accuracy within this dementia cohort was enhanced using the *SynthSeg* algorithm [33], robust to movement artifacts, prior to hippocampal segmentation.

This work also entails several limitations, including the lack of data from pre-dementia onset or from cognitively healthy individuals. Future work should explore if early volume changes observed in HS+ subjects at autopsy are also observed at pre-dementia stages. A modest sample size, unequal number of timepoints per subject as well as a unilateral neuropathological evaluation are other limitations of this study. Our results can serve as foundation for future multi-cohort studies with larger sample sizes to explore volumetric ranges in individuals with HS and AD, and address whether these two can be discerned in pre-mortem MRI.

In summary, we describe reduced hippocampal volumes associated with HS in dementia patients that are detectable earlier than a decade before death, independently of age. HS did not affect the rate of volume decline, and following a comprehensive pathological evaluation, ADNC was the only pathology with a significant effect on hippocampal atrophy rate. These results reveal AD pathology affects hippocampal volume during symptomatic stages of dementia, suggesting its rate of change could be a specific AD biomarker. Our findings also encourage the understanding of HS as a pathology with relevant impact during life, with noticeable early effects in pre-mortem MRI.

## Acknowledgements

The authors thank the participants of the VACS cohort and their family members, as well as the CIEN Foundation staff, especially Paloma Ruiz and Laura Saiz from BT-CIEN and Eva Alfayate from the neuroimaging department. This work has been supported by the Queen Sofia Foundation and the CIEN Foundation. DOC was supported by “la Caixa” Foundation (ID 100010434), with fellowship code LCF/BQ/DR20/11790034. BAS and JEI were supported by a MISTI Global Seed Fund (0000000246). JEI was supported by the ERC (Starting Grant 677697), the NIH (1RF1MH123195, 1R01AG070988, 1UM1MH130981), and ARUK (ARUK-IRG2019A-003).

