## Supplementary Material for "Hippocampal sclerosis of aging at post-mortem is evident on MRI more than a decade prior"

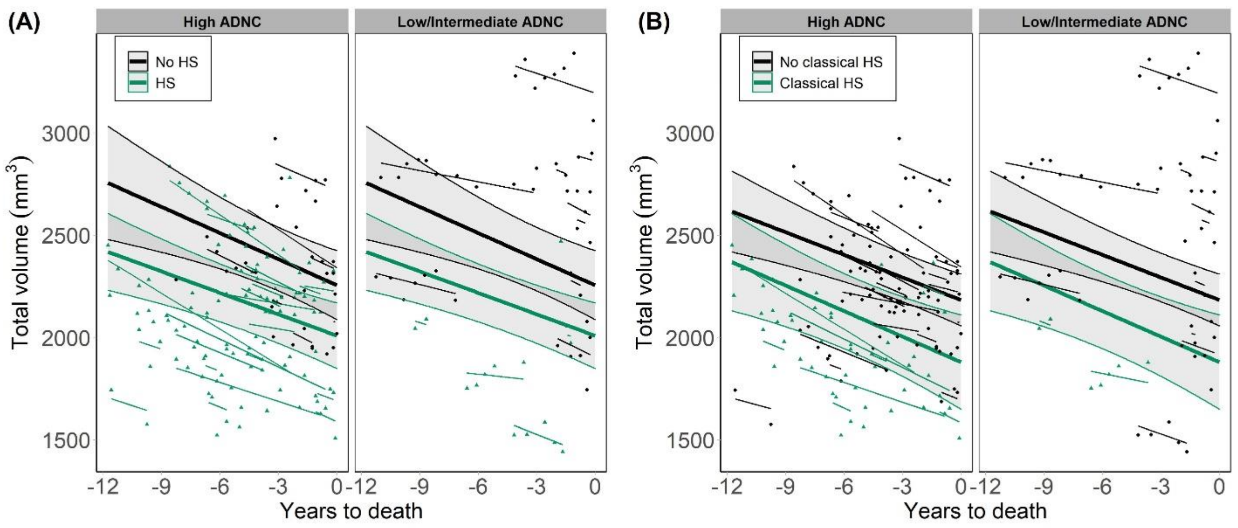


**Figure S1. Hippocampal volume progression as a function of HS and AD burden.** (A) HS groups defined as a function of HS staging, including early and advanced stages. After including AD burden in the statistical model, the effect of HS was still significant ($F=30.8$, $p=.023$), with lower volumes in HS+ compared to HS- patients at 11.75 years before death. (B) HS groups determined by the classical definition of the pathology, including only advanced cases with severe cell loss. Following this classification, the effect of HS was also significant after considering AD burden ($F=26.5$, $p=.01$). ADNC: Alzheimer's disease neuropathological change. HS: hippocampal sclerosis of aging.

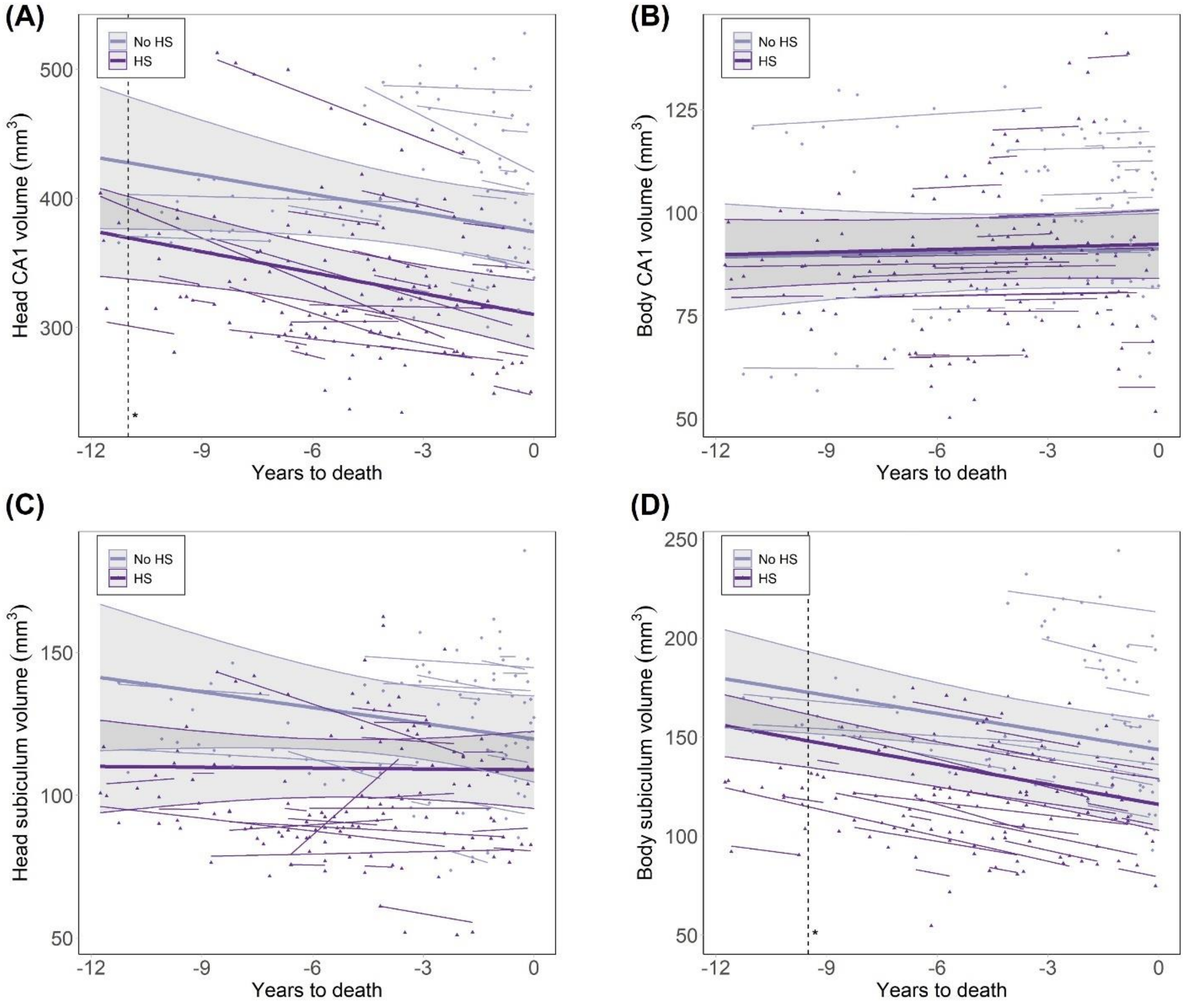


**Figure S2. Volumes of CA1 and subiculum assessed separately as a function of HS.** (A) Comparison of CA1 volumes in the hippocampal head between HS+ and HS- patients. Group classification according to HS staging, including early and advanced stages as HS+. Significantly lower volumes were found in HS+ subjects across the evaluated timespan, with no significantly different slopes between groups. Dotted line indicates the earliest timepoint at which estimated marginal means are significantly different between groups (11 years before death), remaining significant in the interval (-11,0). (B) Volumes of CA1 in hippocampal body, which showed no significant differences between HS+ and HS- groups. (C) Subiculum volumes in the head of the hippocampus, for which no significant differences were found between groups. (D) Volumes of subiculum in the hippocampal body, which were significantly lower in HS+ subjects. Estimated marginal means were significantly different between groups in the interval between 9.5 (dotted line) and 0 years to death. Groups displayed no significant differences in the rate of volume change. HS: hippocampal sclerosis of aging.

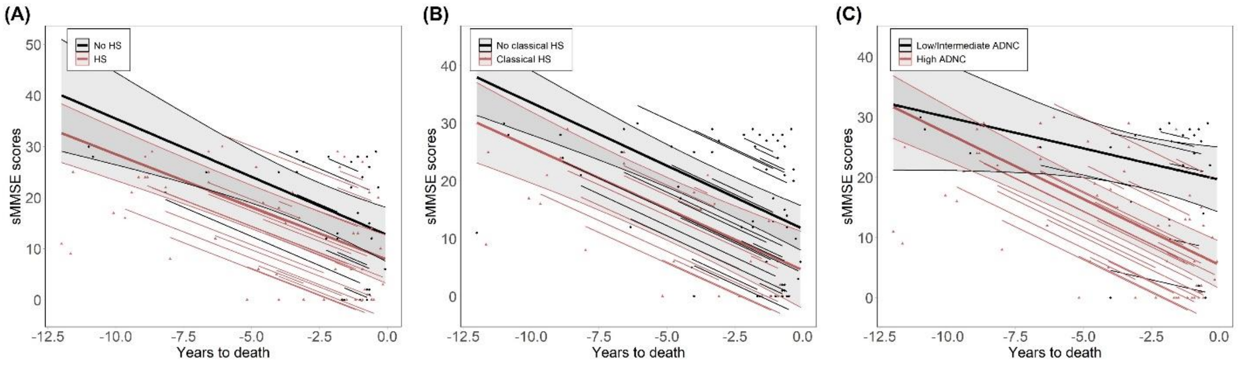


**Figure S3. Progression in severe Mini-Mental State Examination scores as a function of HS and ADNC.** (A) HS groups defined as a function of HS staging showed no significant association with scores across the evaluated timespan. (B) Differences in HS groups following the classical definition of the pathology. The HS+ group showed significantly lower scores ($F=12.2$, $p=.049$), with no differences in rate of decline. (C) Differences between low/intermediate and high ADNC probability groups, the latter showing reduced sMMSE scores ($F=23.1$, $p=2 \cdot 10^{-5}$) and trend-level significantly faster decline ($F=3.8$, $p=.051$). ADNC: Alzheimer's disease neuropathological change. HS: hippocampal sclerosis of aging; sMMSE, severe Mini-Mental State Examination.
