## Supplementary figures and images for "Hippocampal sclerosis of aging at post-mortem is evident on MRI more than a decade prior"

### Supplementary Figure S1

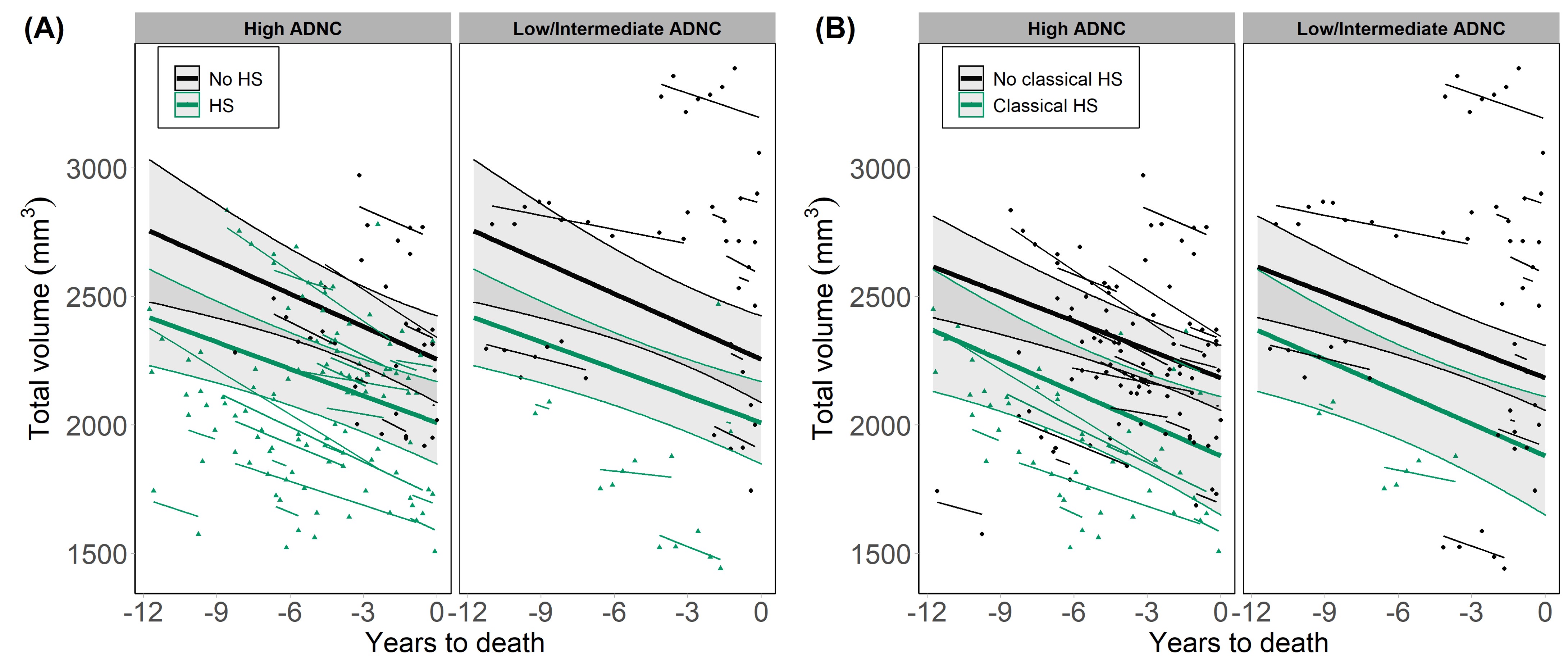

### Supplementary Figure S2

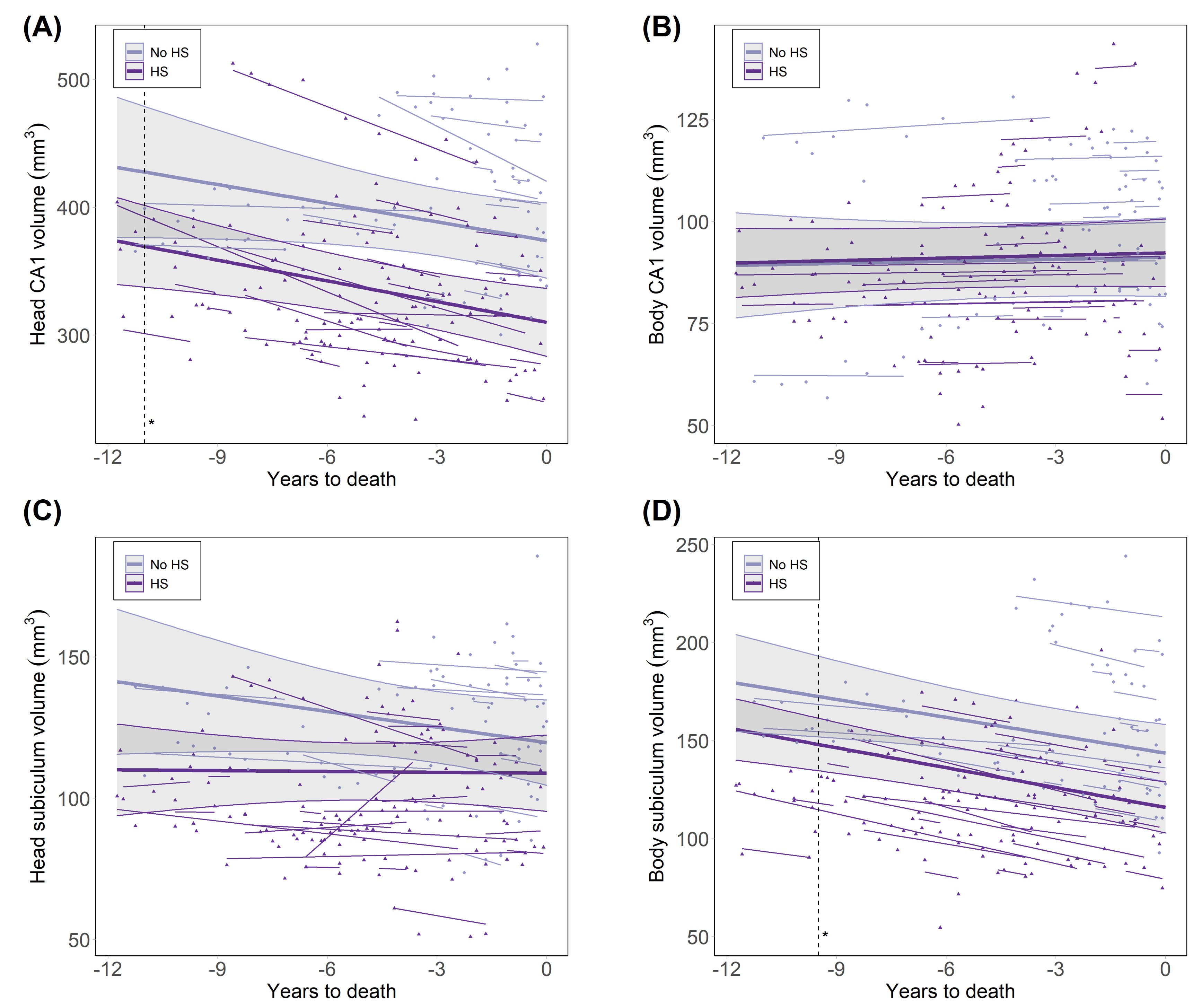

### Supplementary Figure S3

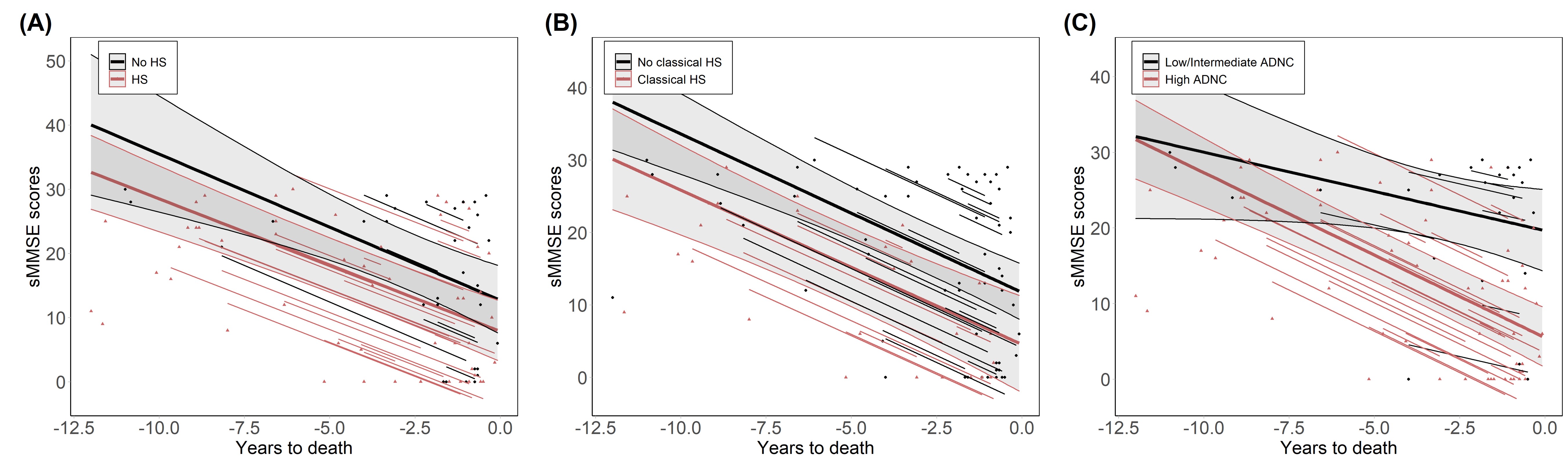
